# Fabrication and Use of a 32-Well LED-Embedded Microplate for Optogenetic Dynamic Control

**DOI:** 10.64898/2026.07.08.737360

**Authors:** Bhavya Jaiswal, Trevor S. Black, Hari R. Namboothiri, Krishna Pochana, Chelsea Y. Hu

## Abstract

**SUMMARY:** This protocol describes how to fabricate, program, and operate a 32-well LED-embedded microplate for optogenetic studies (LEMOS 2.0) inside a microplate reader to enable high-throughput optogenetic stimulation and quantitative gene expression measurements in microbial cultures.

Optogenetic control enables light-actuated regulation of gene expression and provides a programmable interface between living cells and electronic systems. However, routine prototyping of optogenetic constructs remains limited by infrastructure. Existing closed-loop platforms often require chemostats, microfluidics, robotic handling, or custom optical sensors, which can increase cost, reduce accessibility, or constrain measurement performance.

Here, we present LEMOS 2.0, an updated LED-Embedded Microplate for Optogenetic Studies, a low-cost device for optogenetic stimulation and gene-circuit characterization inside standard off-the-shelf microplate readers. LEMOS 2.0 builds on the original LEMOS platform by increasing throughput from 16 to 32 microwells and reducing light leakage between adjacent microwells, allowing dark conditions to be used as an additional illumination state. The device consists of a 3D-printed frame, individually addressable LEDs positioned next to each microwell, a rechargeable battery, and an onboard microcontroller for Bluetooth-based wireless communication. Biocompatible polydimethylsiloxane microwells are cast directly into the device by replica molding, allowing bacterial cultures to be stimulated while optical density and fluorescence are measured by the microplate reader.

This protocol describes the full LEMOS 2.0 workflow, including device fabrication, circuit assembly, Arduino programming, PDMS microwell casting, plate-reader setup, strain and culture preparation, automated experiment execution, device cleanup, and fluorescence/OD_600_ data analysis. As a demonstration, the protocol uses the CcaSR optogenetic system, in which sfGFP expression is activated by green light and repressed by red light. LEMOS 2.0 is intended to make optogenetic perturbation and gene-expression characterization more accessible to wet-lab users, enabling faster design-build-test-learn cycles without requiring specialized bioreactor or microfluidic infrastructure.

## INTRODUCTION

Optogenetics provides a programmable interface between living cells and electronic control systems. By using light to regulate gene expression, cellular behavior can be actuated with temporal precision, reversibility, and minimal perturbation to the culture medium. When paired with fluorescent or luminescent reporters, optogenetic systems allow biological states to be measured electronically and regulated through feedback^1^. This bidirectional optical interface has enabled cybergenetic control of cellular processes, including gene expression, cell growth, and population composition ^2–4^. However, many successful closed-loop optogenetic systems require specialized chemostats^5^, microfluidic devices^4,6,7^, or robotic sampling^2,3,8^, which can limit accessibility and hinder routine design-build-test-learn cycles.

Microplate readers are among the most widely used instruments for measuring microbial growth and gene expression dynamics. They provide reliable optical density and fluorescence measurements across many samples, support environmental control, and are already integrated into standard synthetic biology workflows. However, the illumination paths in standard commercial plate readers are designed for optical measurement, not for programmable, well-specific optogenetic actuation during culture. As a result, optogenetic experiments often require separate illumination devices, manual transfer between stimulation and measurement platforms, or custom readout hardware. These extra steps reduce temporal resolution, introduce handling variability, and complicate closed-loop control.

The LED-Embedded Microplate for Optogenetic Studies, or LEMOS, was previously developed to address this gap ^9^. LEMOS is a microplate-reader-compatible device that contains individually addressable LEDs positioned around culture microwells. The device remains inside a commercial microplate reader throughout the experiment, allowing optical stimulation and fluorescence/OD_600_ measurement to occur in the same platform. In the original implementation, LEMOS supported 16 culture microwells and was used to demonstrate open-loop optogenetic stimulation and closed-loop feedback control of gene expression in batch culture. The device was synchronized with an external computer so that LED illumination could be turned off during measurement windows, minimizing direct interference with fluorescence and optical density readings.

This protocol describes an updated version of the device, LEMOS 2.0 and its use for optogenetic experiments in microbial cultures. The LEMOS 2.0 was redesigned to increase the number of independently addressable culture microwells from 16 to 32, improving experimental throughput and allowing more conditions, replicates, or controller designs to be tested in a single run. The device design was also modified to reduce light leakage between illuminated microwells and neighboring dark microwells, improving the reliability of dark controls and spatially patterned stimulation experiments. These modifications make the platform better suited for comparative optogenetic characterization, multi-condition controller testing, and routine protocol adoption by laboratories that already use microplate readers.

The protocol covers the full LEMOS 2.0 workflow: device fabrication, circuit assembly, Arduino programming, PDMS microwell casting, plate-reader setup, strain and culture preparation, automated experiment execution, device cleanup, and fluorescence/OD_600_ data analysis. Although the example workflow focuses on bacterial optogenetic gene-expression control, the same framework can be adapted to other microbial strains, fluorescent reporters, optogenetic actuator systems, and control algorithms. By combining programmable illumination with standard plate-reader measurements, the updated LEMOS 2.0 platform provides an accessible route for high-throughput optogenetic perturbation and feedback-control experiments in batch culture.

## PROTOCOL

**1. Fabrication of the LEMOS 2.0 Frame and LED Array**

Begin fabricating LEMOS 2.0 by 3D-printing the LEMOS 2.0 frame and placing the LED strips into the device body. The goal is to generate a mechanically stable frame in which each LED is positioned next to the corresponding microwell. Correct LED placement at this stage is important because misalignment can reduce stimulation uniformity and increase optical crosstalk between microwells.

1.1. 3D-print the LEMOS 2.0 inner frame and the casting mold for the PDMS microwells using the provided STL files in the GitHub repository (github.com/synbiosystems/LEMOS-2.0). Remove all support material after printing. We recommend using Polylactic Acid (PLA) filament and a fused deposition modelling (FDM) printer with the following printing parameters: 0.2 mm layer height, 20% infill for the LEMOS 2.0 frame, and 40% infill for the casting mold.

1.2. Cut the LED strips across the center of the soldering pads into 2 sets of 16 LEDs. Each strip end should have three soldering pads available for soldering which is discussed further in Section 2.

1.3. Fold each LED strip in half, with the LEDs facing outward (**Supplementary Figure 1A**).

1.4. Manually conform the LED strip to the device geometry and insert it between the microwells, with the +5V soldering pads oriented downward, noting the arrow on the LED strip (**Supplementary Figure 1A**). Adjust the strip position iteratively until each LED is correctly seated in its slot before continuing.

**2. Assembly and Validation of the Electrical Circuit**

Assemble the power, charging, microcontroller, switch, and LED connections required for wireless device operation. After assembly, the LEMOS 2.0 device should operate from the onboard battery, recharge through the charging module, and receive LED commands from the peripheral Arduino board. Verify each power connection before connecting the LED strips, as incorrect voltage routing can damage the microcontroller or LEDs.

CAUTION: The protocol requires prior proficiency in soldering, including appropriate safety practices (e.g., use of a smoke absorber, protective eyewear, and burn-prevention methods), soldering iron maintenance, and basic through-hole/wire-soldering techniques. Users without this experience should obtain training before attempting the steps in this section.

2.1. Gather all electronic components listed in the Table of Materials and inspect for visible physical damage. Review the wiring diagram provided in the LEMOS 2.0 GitHub repository and keep it available during assembly.

2.2. Configure the step-up voltage converter to output 5V, instead of the default 12V. Following the configuration key on the back of the board, disconnect pads A and B by applying heat from a soldering iron to remove the connection.

2.3. Solder the Arduino Nano BLE Rev2 (referred to hereafter as BLE Nano) VIN pin to the step-up converter OUT + pin and the BLE Nano GND pin to the step-up converter OUT - pin (**Supplementary Figure 1B**).

2.4. Solder the step-up converter IN– pin to the battery charging module negative (-) pin.

2.5. Modify the switch first by removing the mounting pins in each corner. Second, by trimming all pins to 30% of their length. Then, remove completely T1, T3, and B3 (Notation: T = top, B = bottom, numbered left to right from 1 to 4 when looking directly at the pins with the switch on top) (**Supplementary Figure 1D**).

2.6. Separate the two wires of the ‘JST PH 2-Pin Cable – Female Connector, 100 mm’ by pulling the battery-connection ends from the white housing. Trim each to ∼70 mm to better fit in device by trimming from the non-battery-connection ends. (**Supplementary Figure 1C**).

2.7. Solder the red JST cable to B2, leaving the battery-connection end exposed for connecting to the battery’s male connector later (**Supplementary Figure 1D**).

2.8. Solder a connecting wire between B1 and B4.

2.9. Solder the battery charging module B+ pin to B1, being careful to leave the connecting wire from previous step also soldered to B1 (**Supplementary Figure 1E**).

2.10. Solder the step-up converter IN+ pin to T4.

2.11. Solder the battery charging module positive (+) pin to T2.

2.12. Solder the black JST cable to the battery charging module B– pin. Leave battery-connection end loose for connecting to the battery’s male connector later.

2.13. Verify connections under normal device operation mode. Connect Li-ion battery to the circuit via battery connecting wires left loose in soldering steps, with the switch in the center/off position (**Supplementary Figure 1E**).

2.14. Flip the switch to the left/on position (the side with only 2 occupied pins).

2.15. Verify power to BLE Nano via the onboard power LED (labelled ‘ON’, next to the USB port), which should turn green (**Supplementary Figure 1E**).

2.16. Verify power to the step-up converter via its on-board LED, which should turn blue.

2.17. If LED indicators do not illuminate, verify all connections with a multimeter before proceeding.

2.18. Connect the LED strips to the BLE Nano (**Supplementary Figure 1F**) using the 3 soldering pads on the LED strips which are accessible by threading wires through the ports on the side of the frame.

2.19. Solder the Nano GND pin to the LED GND pad using the Nano GND pin closer to the LED strip.

2.20. Solder the Nano D13 pin to the central LED ‘DIN’ pad.

2.21. Solder the Nano VIN pin (also connected to the step-up converter OUT+ pin) to the LED +5 V pad on the closest LED strip end. This requires soldering two wires to the same Arduino pin. One wire solders inside the pin hole, while the other can be soldered to the outside of the pin hole, at the indentation in the Nano board.

2.22. Solder the two LED strips together by soldering connecting wires between the matching pads along the same exterior side used for connecting to the first LED strip.

2.23. Verify connections with a multimeter in continuity mode by touching the VIN pin on the Nano and the +5 V pad on the last unsoldered LED pad. Repeat for GND.

2.24. Secure the LED strips in place using a hot glue gun (**Supplementary Figure 1F**). Move slowly, gluing only one or two LED pairs at a time, allowing the glue to cool between applications. Use tweezers as needed to hold the LEDs in place.

**3. Programming the Central and Peripheral Arduino Boards**

Program the two Arduino boards used for wireless communication and LED control. The central Arduino (Arduino Nano IOT 33) is connected to the computer during an experiment, and it relays commands from the Python script. The peripheral Arduino (Arduino Nano 33 BLE Rev 2) is stationed in the LEMOS 2.0 device and controls the LED array. Both boards must be loaded with the appropriate sketches before starting an experiment.

3.1. Download the Arduino files from the GitHub repository: LEMOS_Central.ino, LEMOS_Peripheral.ino. Download and install the latest version of the Arduino IDE from http://www.arduino.cc/en/software.

3.2. Install the latest version of the board package from the Boards Manager (Tools > Board > Boards Manager) (**Supplementary Figure 2A**).

Central: Arduino SAMD Boards

Peripheral: Arduino Mbed OS Nano Boards

3.3. Install the following Arduino libraries in the Library Manager (Tools > Manage Libraries) (**Supplementary Figure 2B**): ‘ArduinoBLE’ for Bluetooth Low Energy communication between both peripheral and central device, and ‘Adafruit_NeoPixel’ for LED programming.

3.4. Repeat the following sequence once for each Arduino file (LEMOS_Central.ino and LEMOS_Peripheral.ino).

3.5. Open the Arduino file in the Arduino IDE.

3.6. Connect the device to the computer via USB data cable.

3.7. Select the correct COM port for your connected device (Tools > Port). If the COM port is unclear, observe which port appears and disappears when connecting and disconnecting the BLE Nano.

3.8. Select the correct board from the board menu (Tools > Board): Central Device - Arduino Nano 33 IoT, Peripheral Device - Arduino Nano 33 BLE Rev2.

3.9. Upload the script to the device.

A. Optional: To enable a visual indicator of successful connection to the other device, uncomment the listed lines: Central – Lines 34-37, Yellow LED flashes; Peripheral – Lines 99-106, Green LED flashes.

**4. Casting PDMS Microwells by Replica Molding**

Cast optically clear PDMS microwells directly inside the LEMOS 2.0 frame. The PDMS microwells hold the bacterial cultures during plate-reader experiments and must have clear, bubble-free bottoms for reliable OD_600_ and fluorescence measurements. Dust, bubbles, or incomplete curing can introduce well-to-well variation and should be avoided during casting.

CAUTION: Work in a fume hood when handling uncured PDMS. Avoid skin contact with the curing agent.

NOTE: Ensure the fume hood surface is clean before casting, as any dust or debris in the PDMS will compromise its optical transparency and result in inconsistent OD_600_ readings across microwells.

4.1. Sanitize the interior of the LEMOS 2.0 frame where the microwells are cast by spraying 70% ethanol, avoiding direct contact with any electrical components other than the LEDs. Allow the LEMOS 2.0 frame to dry completely before casting PDMS microwells.

NOTE: Ethanol exposure to the LEDs is safe but will damage all other electrical components.

4.2. Spray the casting mold for the PDMS microwells with Ease Release™ 200 and wait 10 minutes for the spray to settle on the plastic surface. A second application after an additional 10 minutes can improve mold release, but it is optional.

NOTE: Ease Release™ 200 will make the casting mold hydrophobic, allowing for easy removal of the casting mold from the cured PDMS microwells.

4.3. Weigh and mix the curing agent and base of SYLGARD™ 184 silicone elastomer (Dow Inc.) to a 10:1 mass ratio of base to curing agent in a disposable plastic cup. We recommend using 2.5 g curing agent + 25 g base for a total of 27.5 g mixture for one LEMOS 2.0 device, scale up proportionally if casting multiple devices simultaneously.

4.4. Mix vigorously for at least 5 minutes using a disposable spatula or wooden stick, scraping the sides and bottom of the cup to ensure complete mixing.

NOTE: Thorough mixing is critical. Undermixed PDMS will not cure uniformly and will yield inconsistent baseline OD_600_ readings across microwells.

4.5. Place the cup containing the PDMS mixture in a vacuum desiccator. Apply vacuum (at least 25 inHg / 85 kPa) for 45–60 minutes, until all visible air bubbles have been removed from the mixture. The PDMS will expand as air bubbles rise to the surface and may overflow its cup. Use a container with at least 3× the volume of the mixture.

NOTE: Residual air bubbles in the cured PDMS will alter the optical transmission of the microwell bottoms and result in inconsistent baseline OD_600_ measurements.

4.6. Secure the LEMOS 2.0 frame to the outer shell using binder clips of appropriate sizes on all four sides.

NOTE: The binder clips minimize the amount of PDMS that seeps into the gap between the outer shell and the LEMOS 2.0 frame. Excess PDMS in this gap can increase the overall device height and prevent the LEMOS 2.0 frame from sitting level within the outer shell. Placing Kapton tape around the bottom perimeter of the LEMOS 2.0 frame and adjacent to the microwells can further reduce seepage.

4.7. Carefully pour the degassed PDMS into the LEMOS 2.0 frame. Use a syringe to add PDMS into each microwell in the frame to ensure they are filled with minimal bubbles.

NOTE: Formation of some small bubbles is unavoidable, and they will eventually rise to the surface and disappear. Pour the degassed PDMS immediately after degassing to minimize bubble reintroduction during handling.

4.8. Insert the 3D-printed casting mold into the PDMS-filled frame. Press the mold gently and evenly, taking care that the circular protrusions in the casting mold sit in the holes in the LEMOS 2.0 frame. The mold defines square microwells aligned next to the LEDs. Wipe away excess PDMS that overflows from the edges with a clean Kimwipe.

4.9. Allow the PDMS to cure at room temperature on a level surface overnight (minimum 16 hours). Elevated temperature curing (e.g., 37 °C for 4 hours) may be used to accelerate curing if needed; confirm all bubbles have been removed from the PDMS before doing so.

4.10. Once cured, carefully remove the casting mold by gripping it at the edges and pulling it up with a steady force. The PDMS microwells should remain in the frame. Inspect each microwell to ensure that the walls are optically clear and the bottom is free of bubbles.

4.11. The LEMOS 2.0 device with fresh PDMS microwells is now ready for use.

**5. Configuring the Microplate Reader and Data Export Workflow**

Configure the microplate reader for LEMOS 2.0-compatible kinetic measurements and automated data export. The reader must recognize the custom LEMOS 2.0 well layout, acquire OD_600_ and fluorescence measurements at defined intervals, and export data files to the folder monitored by the Python control script. Perform a baseline measurement with sterile medium before biological experiments to confirm that the PDMS microwells and plate-reader settings produce consistent readings. This protocol uses a BioTek Synergy H1 microplate reader operated via BioTek Gen5 software v3.14.

5.1. Launch Gen5.

5.2. Go to Plate Types and import LEMOS 2.0.xml from the GitHub or alternatively set up the custom plate layout for LEMOS 2.0 in Gen5 by referring to Supplementary Protocols Section 1.

5.3. Perform a baseline measurement with the LEMOS 2.0 device (microwells filled with 200 µL of sterile M9CA media, covered with Breathe-Easy film). Record the OD_600_ and fluorescence baseline values for all 32 microwells and compare these values to those obtained from a standard 96-well microplate. If a large discrepancy is observed, re-cast the PDMS microwells, as bubbles or debris in the cured PDMS are the most likely cause.

5.4. Open the experiment protocol from the GitHub repository in Gen5.

NOTE: A pre-configured experiment file is available in the GitHub repository for convenience. However, the settings are described in detail in Supplementary Protocols Section 2 to provide a reference for troubleshooting for users with different versions of Gen5.

**6. Preparation of Bacterial Cultures for LEMOS 2.0 Experiments**

Prepare bacterial cultures for a LEMOS 2.0 optogenetic time-course experiment. A cautious approach to culture preparation is important because starting density and pre-experiment light exposure can affect the measured gene-expression dynamics. The demonstration strain is *Escherichia coli* (*E. coli*) MG1655 carrying pSR58.6 and pNO286-3 ^10^. pSR58.6 contains PcpcG2-sfGFP and constitutive CcaR, chloramphenicol resistance, and a pColE1 origin. pNO286-3 contains constitutive CcaS, spectinomycin resistance, and a p15A origin. Together, these plasmids reconstitute a green-light-inducible, red-light-repressible gene expression system.

6.1. Inoculate a single colony of the CcaSR optogenetic strain and a fluorescence-negative control strain into separate 14 mL round-bottom culture tubes, each containing 2 mL of M9CA broth supplemented with 50 µg/mL spectinomycin and 25 µg/mL chloramphenicol. Use this media formulation for all subsequent culture steps throughout the experiment.

6.2. Cover each tube with aluminum foil and incubate overnight (14–16 hours) at 37 °C with shaking at 220 rpm.

NOTE: Foil wrapping maintains dark conditions and prevents unintended activation of the CcaSR system. Alternatively, cultures can be incubated under constant red light by housing the tubes in an LED enclosure wired with the same electrical setup described in section 2, which more closely mimics the repressed state of the system prior to the experiment.

6.3. Transfer 100 µL of each overnight culture into 2 mL of fresh media in a new foil-wrapped culture tube. Incubate for 4 hours at 37 °C with shaking at 220 rpm.

6.4. After 4 hours, measure the OD_600_ of each sub-culture using the microplate reader. Cultures should have reached an OD_600_ of 0.4–0.6.

6.5. Dilute each culture to OD_600_ = 0.1 in fresh media, pipette 200 µL into each microwell of the LEMOS 2.0 device taking care while pipetting to avoid the formation of air bubbles, and seal the device with Breathe-Easy film.

NOTE: Breathe-Easy film is gas-permeable, maintaining aerobic conditions for cell growth, while remaining optically transparent to avoid interference with fluorescence and OD_600_ measurements.

**7. Initiation of a LEMOS 2.0 Optogenetic Time-Course Experiment**

Start the automated LEMOS 2.0 experiment by coordinating the microplate reader, the central Arduino, and the Python control script. During the run, the computer coordinates plate-reader measurements, data export, Bluetooth communication, and LED actuation. It is critical that the Gen5 window is visible because the Python script uses screen-based automation to interact with the plate-reader software. A successful start is indicated by exported data files appearing in the expected folder, terminal output confirming Bluetooth communication, and LED commands updating at each control interval.

7.1. Slide the switch to the left to power on the LEMOS 2.0 device. The first LED on the LED strip in LEMOS 2.0 will illuminate green.

7.2. Connect the central Arduino to the computer via USB and select it in the Arduino IDE from the drop-down menu (Tools > Board).

7.3. Open the Serial Monitor and connect the peripheral Arduino staged in LEMOS 2.0 to the central Arduino.

7.4. Turn off the indicator LED on the LED strip by entering the letter ‘o’ into the serial monitor.

7.5. Open the experiment file LEMOS-start-data.xpt in Gen5 and take a baseline reading of the cultures in the device, ensuring that the OD_600_ and fluorescence values are consistent with expectations. Close experiment file.

7.6. Open the protocol file saved in step 5.4 in Gen5 and save as an experiment by referring to Supplementary Protocols Section 3.

7.7. In a suitable Integrated Development Environment (IDE), open ble_control.py. We recommend using Visual Studio Code (VS Code).

NOTE: Ensure that the working directory is set to the folder which contains all the Python scripts, so they can be imported during script execution.

7.8. Scroll down to the Main section in ble_control.py. Change the file variable to match the filename chosen for the Gen5 experiment file in step 7.6.

7.9. Setup the Gen5 experiment file and Python script side by side as shown in Figure 4.

7.10. Click **Run** on the Gen5 experiment, then abort it once the kinetic step begins.

NOTE: The Python script looks for a **Continue** button after clicking **Run**. However, in a fresh experiment file, this button is not displayed until the second reading is taken or the kinetic run is aborted.

7.11. Run the Python script. Do not minimize or cover the Gen5 window during the run.

NOTE: The Python script uses screen-based automation and must be able to locate the Gen5 window approximately every 20 minutes to handle data export after each kinetic cycle. Obscuring or minimizing the window during this time may disrupt the experiment.

**8. Device Cleaning, PDMS Removal, and Battery Recharging**

Clean and reset the LEMOS 2.0 device after the experiment. Remove the bacterial cultures and used PDMS microwells, sanitize the reusable frame and shell, and recharge the battery for the next run. Avoid exposing the non-LED electronic components to ethanol or excessive mechanical force during cleaning.

8.1. Gently peel the Breathe-Easy film from the top of the LEMOS 2.0 device and discard.

8.2. Using a pipette, aspirate and discard all cell suspension fluid from each microwell according to biological waste disposal procedures.

8.3. Separate the outer shell from the LEMOS 2.0 frame by applying steady, even force. Avoid excessive bending or twisting of the PLA frame, as this may cause cracking or permanent deformation.

8.4. Peel away the bulk PDMS layer from the exterior of the frame and use tweezers to carefully remove PDMS microwells from the LEMOS 2.0 frame.

8.5. Sanitize the LEMOS 2.0 frame and outer shell with 70% ethanol, avoiding direct contact with any electrical components other than the LEDs, as described in step 4.1.

8.6. Recharge the Li-ion battery for the next experiment: 1) Slide switch to the right/charging position, 2) Connect USB to charging module, 3) Confirm charging via the onboard LED which will flash while charging (**Supplementary Figure 1E**).

**9. Analyzing Fluorescence and Growth Dynamics**

Process the fluorescence and OD_600_ data generated during the LEMOS 2.0 experiment. The analysis notebook imports the exported plate-reader data, applies blank and negative-control corrections, calculates normalized fluorescence, and generates time-course plots. Before running the notebook, confirm that the well assignments and condition labels match the experimental layout.

9.1. Open FL_OD_Data_Analysis_Python.ipynb.

NOTE: This uses fluorescence (FL) and optical density (OD) data, which are stored in these files, generated during the run: Datafile/fl.csv and Datafile/od.csv.

9.2. Set the conditions variable with condition names and wells.

9.3. Set neg_wells equal to a bracketed list of negative control well names.

9.4. Set od_blank and fl_blank to averages across blank wells, or to known values if no blank was run.

9.5. Set colors_4_figures to desired colors (full list of named colors available at https://matplotlib.org/stable/gallery/color/named_colors.html).

9.6. Set figsize to desired width and height of resultant figures in inches.

9.7. Confirm filenames listed under “Constants”.

9.8. Run notebook. Figures are generated as .png files.

**10. Troubleshooting**

10.1. When switch is turned on, Arduino Nano’s power LED does not illuminate.

A. First, confirm the battery is charged by connecting the device to a USB power source in charging mode; the indicator LED should flash green in a few minutes. If the battery was drained, allow it to charge fully before retrying.
B. If the battery is charged and the LED still does not illuminate, use a multimeter to measure the voltage across the BLE Nano’s VIN and GND pins.

i. 0 V: A wiring connection is broken. Inspect each solder joint along the battery → switch → step-up converter → Nano path and re-flow any joint that looks cold or incomplete.
ii. 5 V: Power is reaching the Nano correctly, so the board itself may be damaged. Attempt to power the Nano via cable to confirm this. If power LED still does not illuminate, a new Nano must be soldered in to replace the faulty Nano.
iii. Greater than 5 V: The step-up converter has not been configured to output 5 V. Confirm pads A and B have been disconnected on the step-up converter, as described in the device assembly section.

10.2. Python script fails to run.

A. Confirm the script is opened from within its full project folder rather than as a standalone file — ble_control.py depends on other local modules and will fail if opened independently.
B. Confirm that Arduino IDE is closed, since the Arduino can only have one active connection at a time. The operating system grants access to whichever program opens the port first.
C. Confirm that the Gen5 protocol is set up correctly, especially if using different software. Refer to Supplementary Protocols Section 2.

10.3. Experiment stops unexpectedly.

A. Stops after the first cycle: The Gen5 data export path is likely misconfigured. Confirm the export destination matches the folder the Python script expects to read from.
B. Stops after several cycles: pyautogui occasionally fails to recognize on-screen elements, often after a change in screen resolution, display scaling, or window size. Re-capture the reference screenshots used by the script and keep the Gen5 window active and in the foreground throughout the run; the Python IDE window can be minimized.
C. In the event that Gen5 reports that the chamber temperature is outside the target range, allow the reader to equilibrate before continuing. Do not override the temperature warning, because doing so can trigger an audit report window that must be closed manually and may interrupt automation.

10.4. Unusually high or inconsistent OD600/fluorescence baseline values across microwells.

A. Inspect the PDMS microwells for trapped dust or debris, which scatters light and inflates readings. Avoid leaving PDMS uncovered and exposed to ambient air for extended periods before casting or use.

10.5. Cells show no detectable response to light despite running the experiment.

A. Confirm LED “on” commands are being issued by checking both the IDE terminal output and the BLE communication log.
B. If commands are being sent but no light is visible, check the LED soldering connections with a multimeter.

## REPRESENTATIVE RESULTS

LEMOS 2.0 is a 32-well, microplate-reader-compatible platform^11^ for programmable optogenetic stimulation and quantitative gene-expression measurement (Figure 1A,B). Each microwell is paired with an individually addressable WS2812B light-emitting diode (LED), controlled by an Arduino Nano 33 BLE Sense Rev2 through the NeoPixel protocol. The device communicates with an external computer through Bluetooth Low Energy, allowing illumination commands to be updated during the experiment. Fresh polydimethylsiloxane (PDMS) microwells are cast into the device before use by replica molding (Figure 1C,D). E. coli cultures grown in LEMOS 2.0 at 30 °C under constant LED illumination showed reproducible growth across microwells, indicating that the device supports bacterial culture and plate-reader measurement over the experimental time course^9^ (Figure 1E).

**Figure 1.**
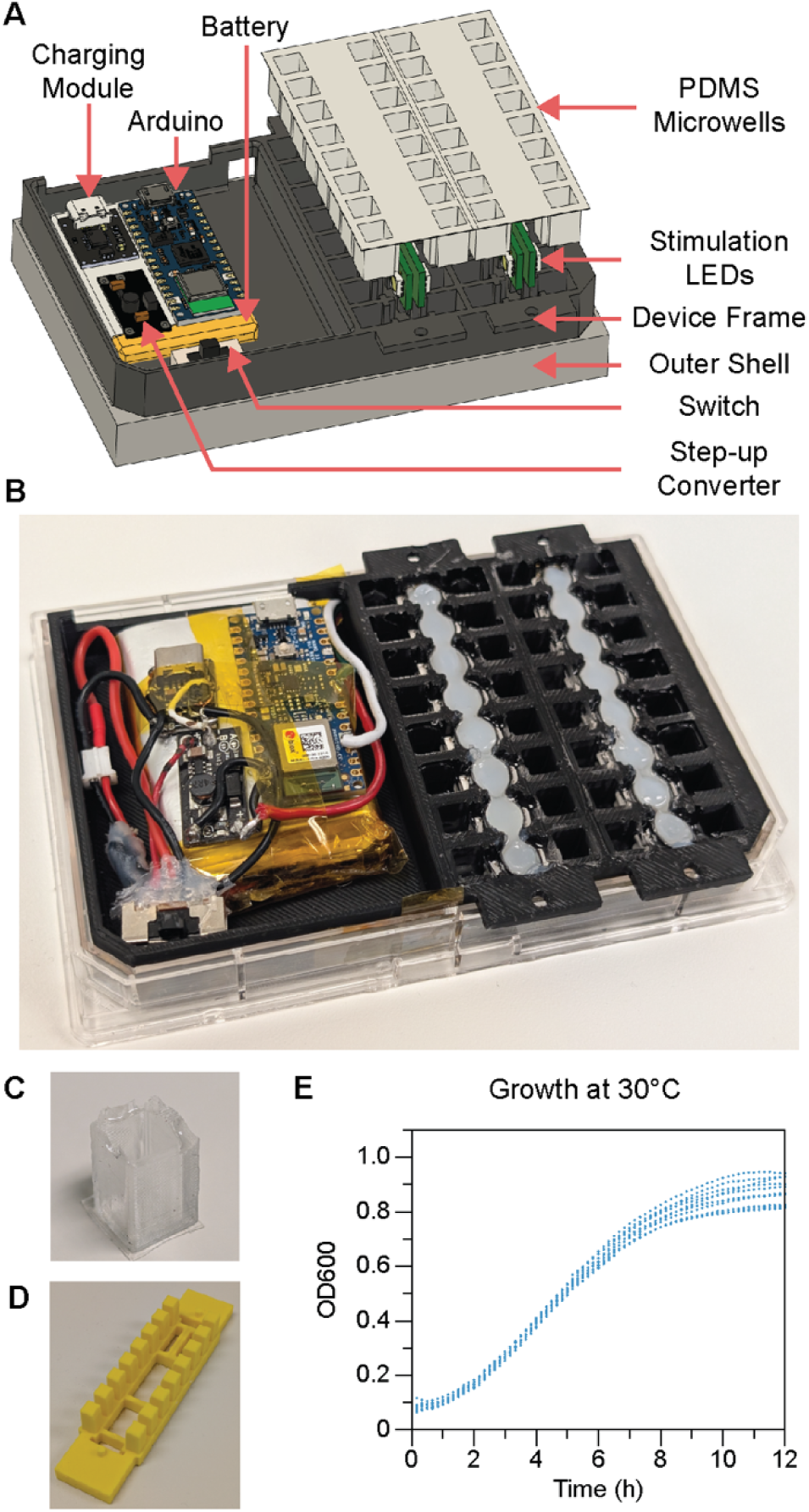
LEMOS device construction and cell growth characterization. (A) Exploded schematic of the LEMOS device showing PDMS microwells, stimulation LEDs, battery, switch, microcontroller, charging module, device frame, and outer shell. (B) Assembled device with PDMS microwells filled with deionized water. (C) PDMS microwell cast in the LEMOS device. Reproduced from Namboothiri et al., *ACS Synthetic Biology* (2026), licensed under CC BY 4.0. (D) 3D-printed mold for casting microwells (E) Growth curves of *E. coli* in LEMOS at 30 °C (N = 24 technical replicates). Gray shading marks the logarithmic (exponential) growth phase. N denotes the number of technical replicates.

To demonstrate optogenetic gene-expression control, we used the CcaSR v3.0 system^12^, a green-light-activated and red-light-repressed two-component system that controls sfGFP expression (Figure 2A). Under green light, CcaS activates CcaR, which drives PcpcG2-dependent sfGFP expression. Under red light, CcaS promotes CcaR dephosphorylation, reducing PcpcG2 activity and repressing sfGFP expression. The CcaSR plasmids were transformed into E. coli MG1655 and used for the experiments described in this protocol (Figure 2B). During the experiment, the microplate reader measured OD_600_ and fluorescence every 10 min. To prevent LED illumination from interfering with optical measurements, the LEDs were programmed to turn off during measurement windows at the beginning and end of each 10-min interval (Figure 2C).

**Figure 2.**
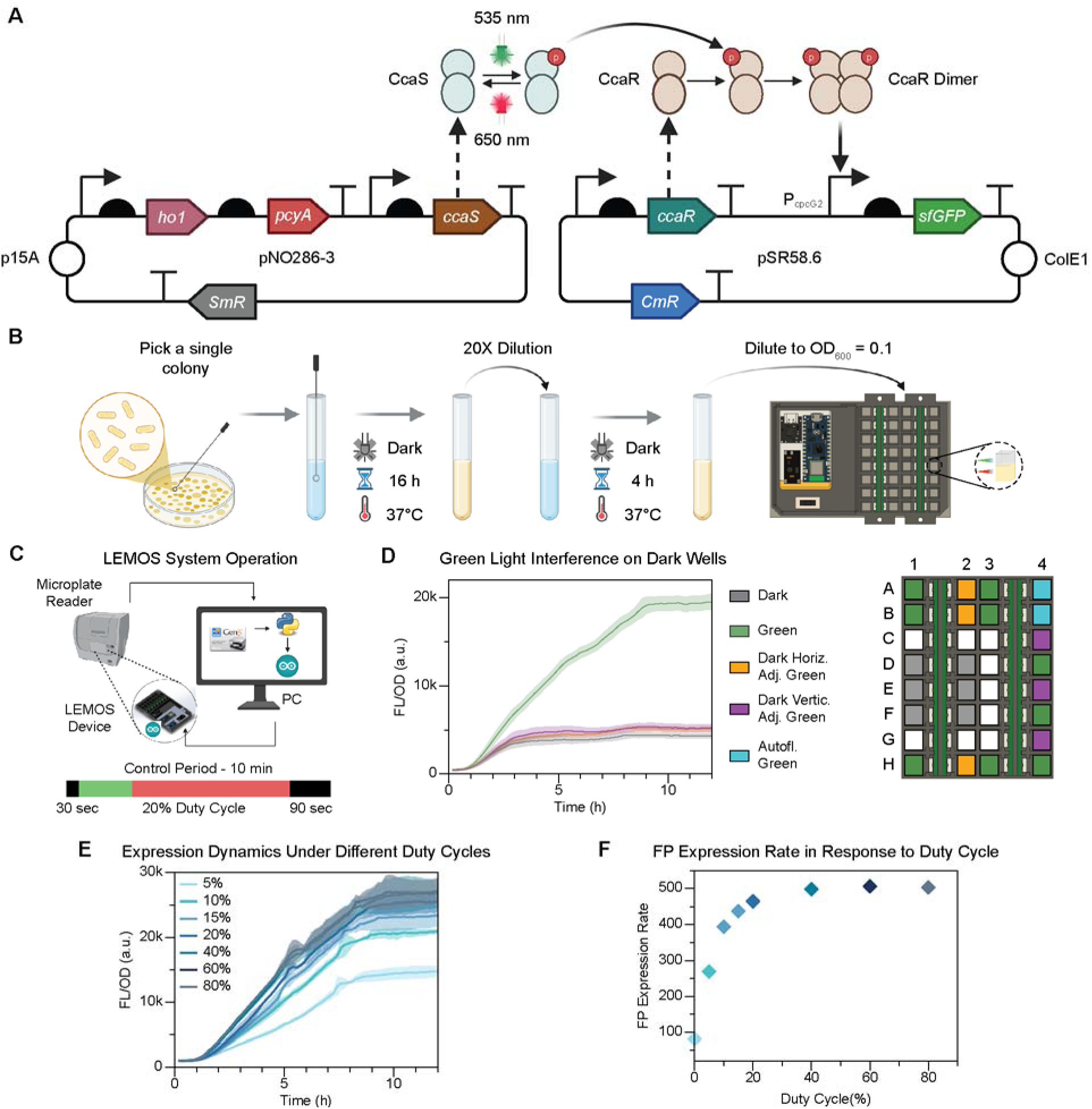
Optogenetic system characterization in LEMOS and open-loop control dynamics. (A) Plasmid map of the CcaSR two-component optogenetic system: green light (522 nm) activates CcaS/CcaR to upregulate sfGFP expression, and red light (620 nm) downregulates the expression. (B) Protocol for carrying out optogenetic experiments with CcaSR Optogenetic system. (C) LEMOS operation schematic. During the 12-h time course, the device remains in the microplate reader. The reader measures sfGFP fluorescence and OD_600_ and streams data to a computer, which handles timekeeping and operates Arduino routines that command the onboard microcontroller to temporarily disable LED illumination during each measurement. Adapted from Namboothiri et al., *ACS Synthetic Biology* (2026), licensed under CC BY 4.0. (D) Effect of green-light interference on unstimulated cells in the dark condition. In the plate layout, green boxes indicate green-light microwells (N = 9), grey boxes indicate microwells in the dark condition with no light interfering in the surrounding microwells (N = 6), purple boxes indicate dark microwells (N = 3) vertically adjacent to green-light microwells, yellow boxes indicate dark microwells (N = 3) horizontally adjacent to green-light microwells and blue boxes indicate green-light microwells with a control strain to measure autofluorescence (N = 2). (E) Open-loop responses in LEMOS to varying green-light duty cycles (5–80%, N = 3) as actuating parameter. Solid lines indicate means; shaded bands indicate standard deviations. (F) Average rate of expression during the exponential phase of growth with varying duty cycle. N denotes the number of technical replicates.

We first evaluated interwell crosstalk by placing dark-condition microwells vertically or horizontally adjacent to microwells under constant green-light illumination (Figure 2D). Dark microwells adjacent to constant green-light microwells showed no significant increase in expression relative to dark microwells without illuminated neighboring microwells, indicating minimal light leakage between microwells. Based on these results, a PWM value of 1 out of 255 was used for subsequent experiments to provide sufficient optical stimulation while minimizing interwell crosstalk.

Open-loop stimulation was then used to define the actuating parameter for LEMOS-based optogenetic control. The actuating parameter was the green-light duty cycle, defined as the fraction of each active illumination window assigned to green light, with the remaining illumination time assigned to red light. Each control interval lasted 10 min, with LEDs active for 8 min and off for the remaining 2 min to avoid interference with plate-reader measurements. Green-light duty cycles from 5% to 80% produced graded sfGFP expression, showing that duty cycle can be used as a tunable input for optogenetic regulation (Figure 2E). The expression rate increased as the duty cycle rose from 5% to 40%, with no significant additional increase at higher duty cycles (Figure 2F), suggesting that the system approaches saturation under stronger green-light stimulation.

Finally, we implemented closed-loop feedback control to test whether LEMOS 2.0 could dynamically regulate gene expression toward a defined set point (Figure 3A). sfGFP fluorescence normalized by OD_600_ served as the measured output, the external computer acted as the controller, and the LEMOS 2.0 LEDs served as the actuator. The controller compared the measured FL/OD_600_ value with the assigned set point and updated the green-light duty cycle for the next 10-min interval. We implemented proportional-integral-derivative (PID) control, where the duration of green light exposure in each cycle, *t_green_*, was computed as:

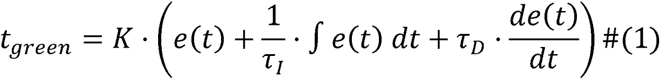

**Figure 3.**
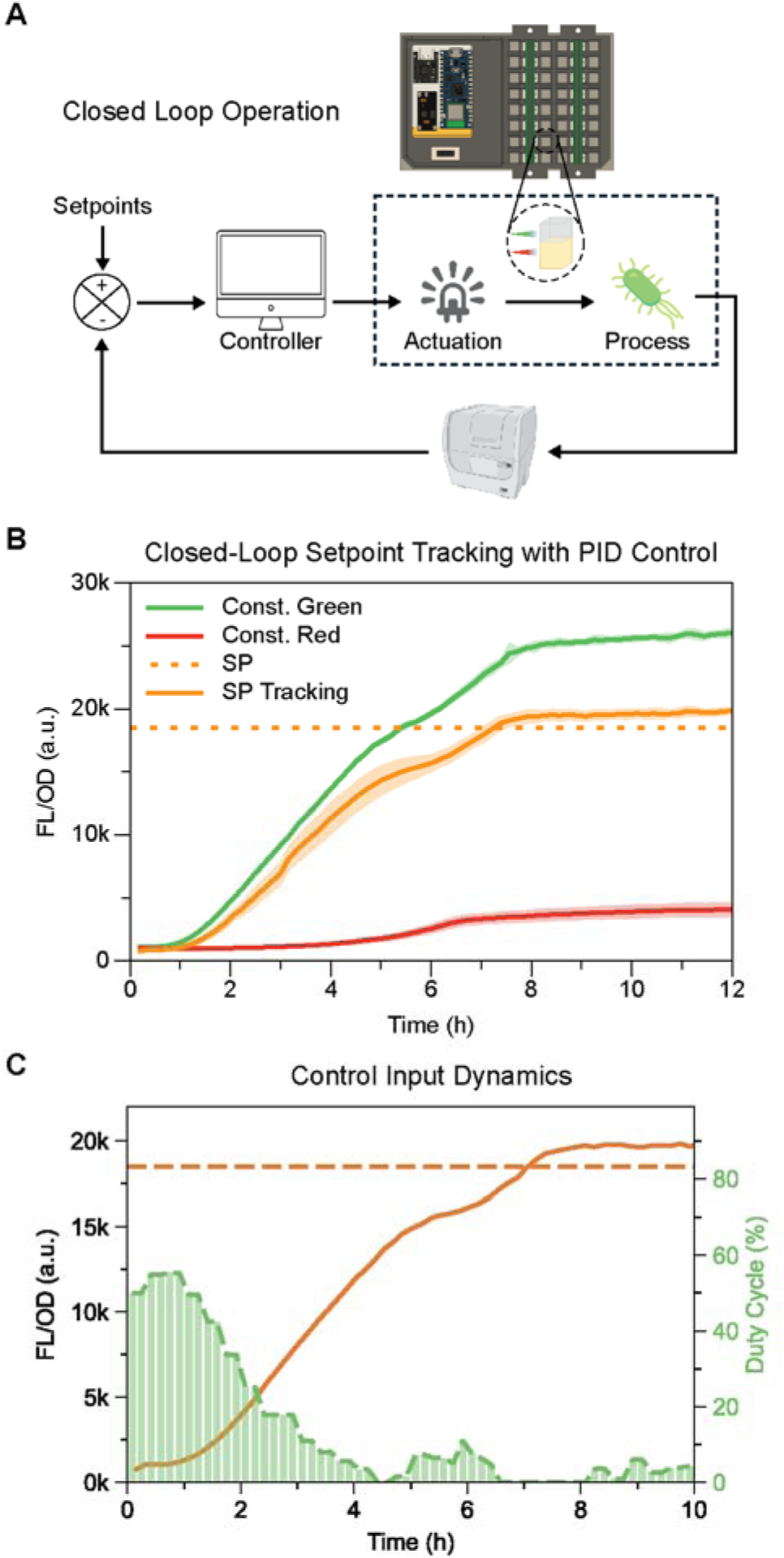
Closed-loop feedback control in LEMOS 2.0. (A) Schematic of closed-loop operation: the microplate reader functions as the sensor that measures FL/OD_600_, the computer functions as the controller, and the LEMOS 2.0 device actuates LEDs to regulate gene expression. The process plant is gene expression in living bacterial cells. Adapted from Namboothiri et al., *ACS Synthetic Biology* (2026), licensed under CC BY 4.0. (B) Proportional Integral Derivative (PID) control tracking of FL/ OD_600_ set point (SP = 18.5 × 10^5^ a.u.). Constant-light references are green and red (N = 3). The set point tracking runs is described by orange lines (N = 3). Solid lines indicate the mean FL/OD_600_; shaded bands indicate standard deviations. Dotted line marks the set point. (C) Representative duty-cycle commands generated by the PID controller during closed-loop setpoint tracking. Bar height indicates the fraction of green light delivered during each illumination interval. N denotes the number of technical replicates.

**Figure 4.**
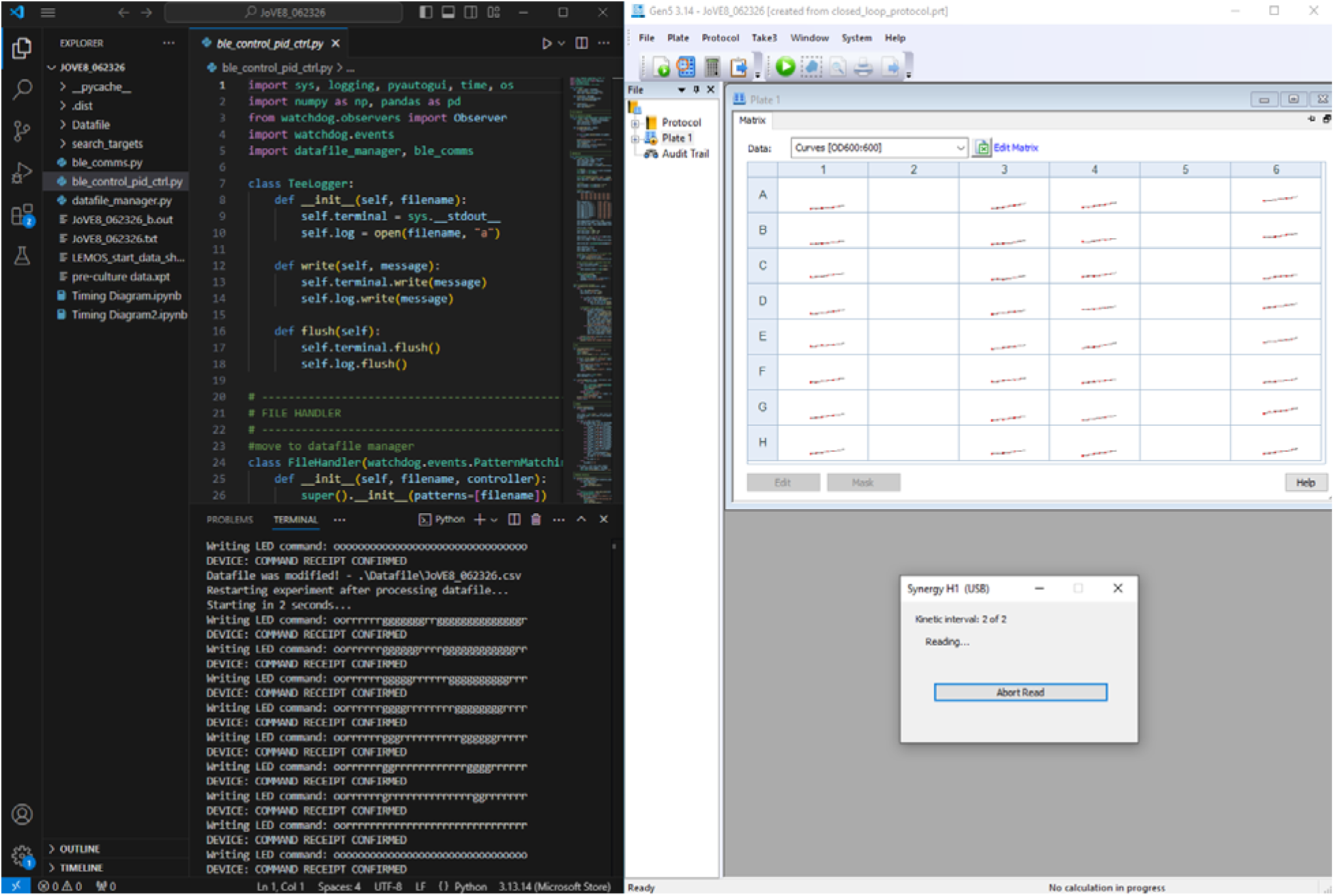
LEMOS 2.0 software setup during an experiment. Sample screenshot showing the Python control script running in VS Code (left), with LED commands populating in the terminal as they are sent to the Arduino microcontroller. The project directory containing the main script and associated function modules is visible in the file explorer panel. On the right, the plate reader software shows a kinetic measurement cycle in progress; at the end of each cycle, the Python script automatically triggers the next run via the Gen5 interface and updates the LED duty cycle commands accordingly.

Here, *e*(*t*) is the error between the measured FL/OD_600_ signal and the set point at time t. *K* is the proportional gain, *τ_I_* is the integral time constant, and *τ_D_* is the derivative time constant. The closed-loop control parameters were selected using the Gene Expression Across Growth Stages (GEAGS) model previously reported by our group^13^. Using these parameters, the measured signal approached and tracked the target FL/OD_600_ set point (Figure 3B).

The corresponding duty-cycle commands are shown in Figure 3C. Bar height represents the fraction of green light delivered during each 8-min illumination window. The duty-cycle command changed in discrete steps because the controller updated the input once every 10 min. As sfGFP expression approached the set point, green-light exposure decreased and red-light exposure increased. After the measured signal reached the set point, a low level of green-light input was still observed. This residual control input likely reflects the derivative term responding to the sharp change in expression rate near the set point. These results show that LEMOS 2.0 can implement model-guided PID feedback control of optogenetic gene expression in batch culture.

## DISCUSSION

This protocol describes the fabrication and operation of LEMOS 2.0, a 32-well LED-embedded microplate for optogenetic stimulation and feedback-control experiments inside a standard microplate reader. The representative experiments show that LEMOS 2.0 supports microbial growth, reduces light leakage into dark-condition microwells, enables duty-cycle-based optogenetic actuation, and implements closed-loop PID control of gene expression in batch culture.

Several steps are critical for reliable performance. During fabrication, each LED must be aligned with the corresponding microwell to maintain stimulation uniformity and reduce optical crosstalk. During circuit assembly, the step-up converter must be configured to 5 V before connecting the microcontroller and LED strips. PDMS casting also requires care, because bubbles, dust, debris, or incomplete curing can alter optical transmission through the microwell bottoms and produce inconsistent OD_600_ or fluorescence baselines. A baseline measurement with sterile medium should therefore be performed before biological experiments.

Software configuration is another common source of failed experiments. The current workflow depends on coordinated operation of the plate-reader method, Gen5 data export, the Python control script, and Bluetooth communication between the central and peripheral Arduino boards. The export path and file format must match the structure expected by the Python script, and the Gen5 window must remain visible during the run because the automation uses screen-based interactions. Users adapting the workflow to another plate-reader model or software package should preserve the same timing, measurement, export, and parsing logic while modifying the automation layer as needed.

LEMOS 2.0 is currently optimized for batch-culture experiments. Because growth rate, resource availability, and gene-expression dynamics change during batch culture, LED intensity, duty-cycle range, sampling interval, and controller parameters may need to be recalibrated for other strains, reporters, optogenetic systems, or growth conditions. Future versions could incorporate in-plate channels or reservoirs for controlled medium exchange and dilution, allowing longer experiments while maintaining more consistent growth conditions. Additional designs could also support alternative LED wavelengths or reduce dependence on screen-based software automation.

## ACKNOWLEDGMENTS

The authors B.J., T.S.B and H.R.N. were supported by the Texas A&M Engineering Experiment Station.

## MATERIALS

Table of Materials used can be found in the SI.

