## Supplementary material for "Fabrication and Use of a 32-Well LED-Embedded Microplate for Optogenetic Dynamic Control": SI file

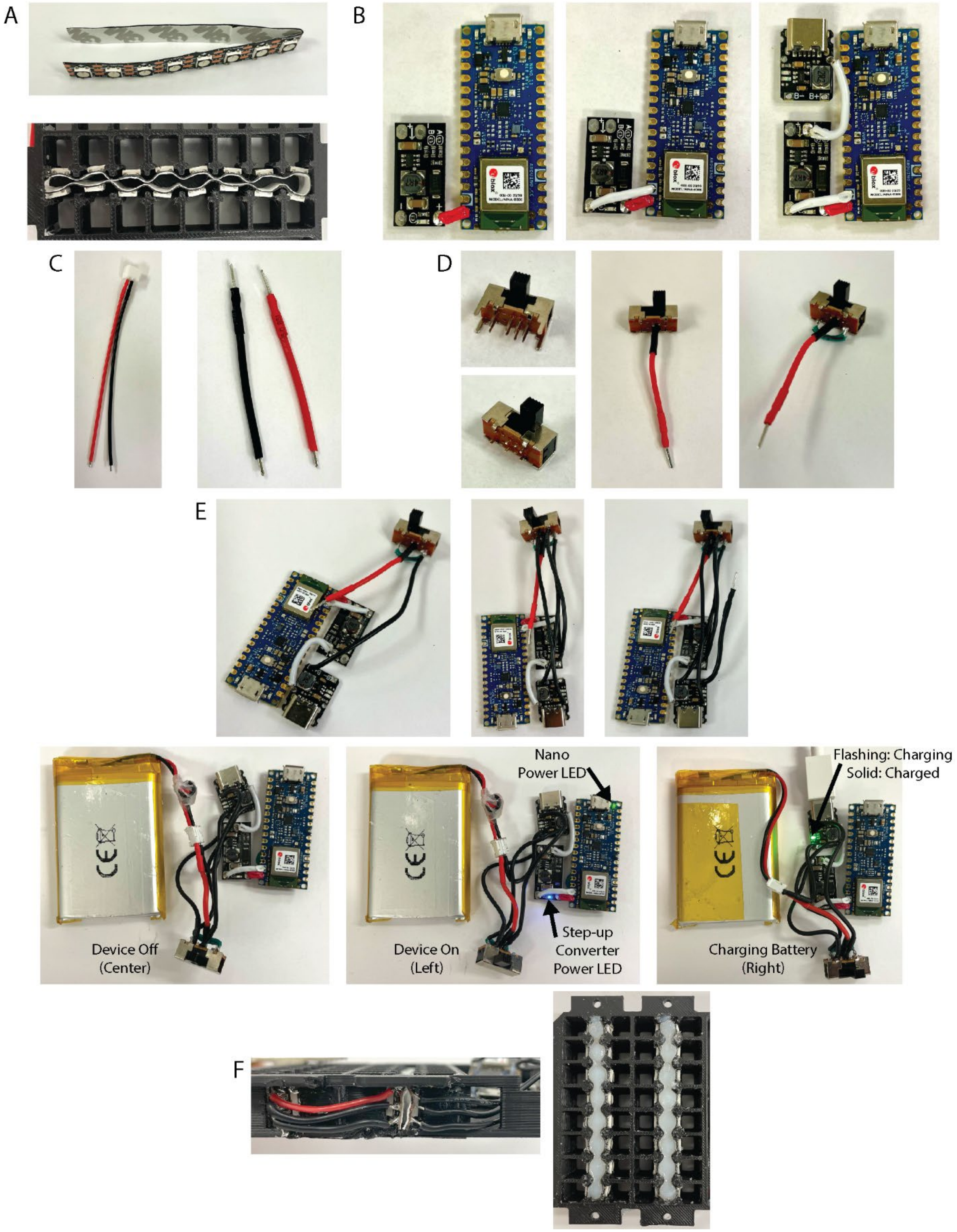


**Figure S1. Assembly Images.** (A) Bent LED strip and molded strip inside the device. (B) Nano BLE Sense Rev 2 solder to step-up converter, which is soldered to the battery charging module. (C) Original and modified versions of the 'JST PH 2-Pin Cable – Female Connector 100mm'. (D) Original switch, modified switch, soldered red JST PH cable, and short jumper soldered between two pins on either side of the bottom row. (E) Incorporation of the switch into the circuit assembly and powering the circuit. (F) Soldering to the LED strips and hot gluing the two LED strips in place.

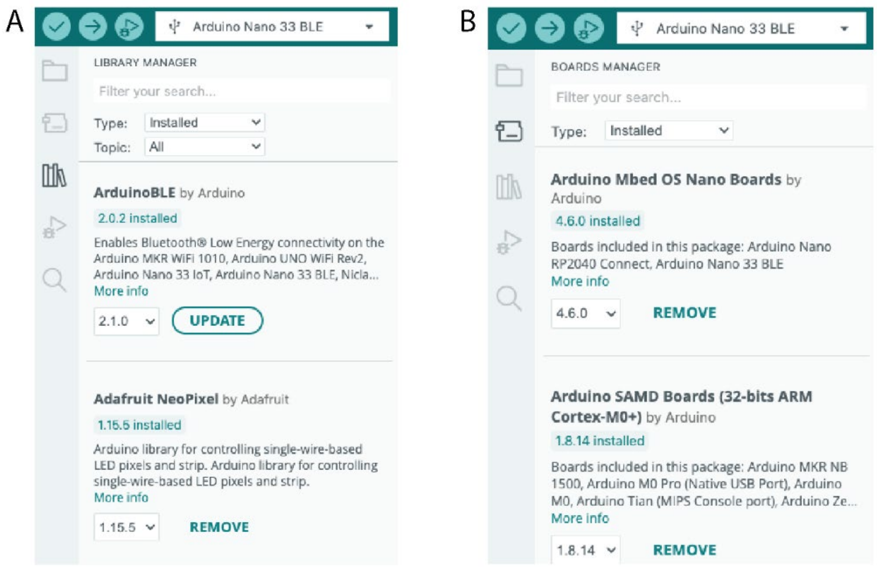


**Figure S2. Arduino IDE Setup Screenshots.** (A) Boards Manager screenshot, showing a search for one of the required boards. (B) Library Manager screenshot, showing two of the required libraries.

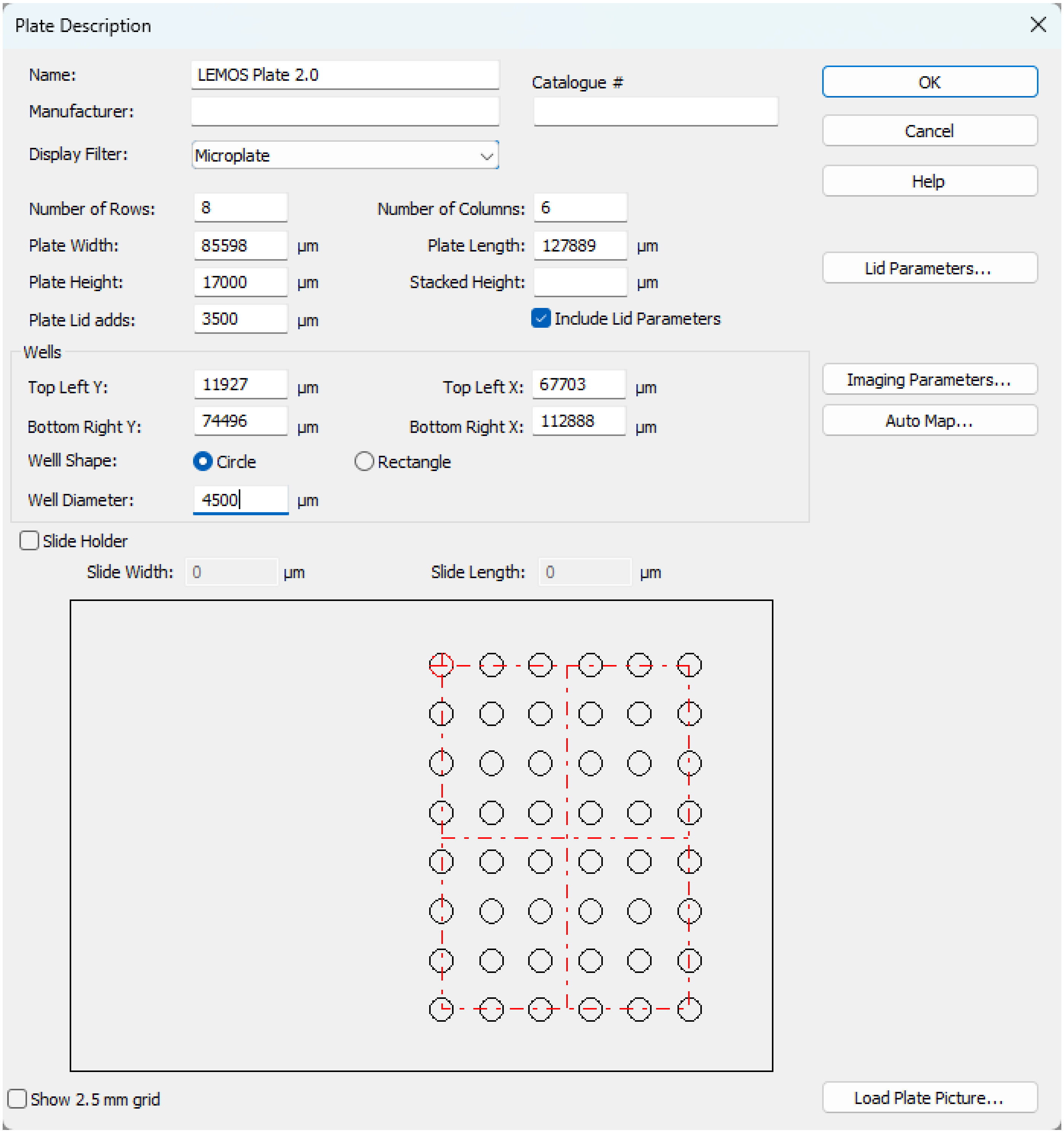


**Figure S3. LEMOS Layout – Plate Description.**

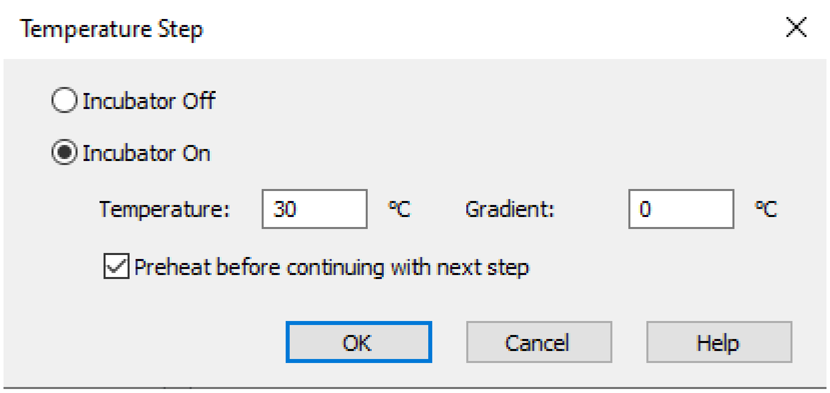


**Figure S4. LEMOS Protocol – Temperature Step.**

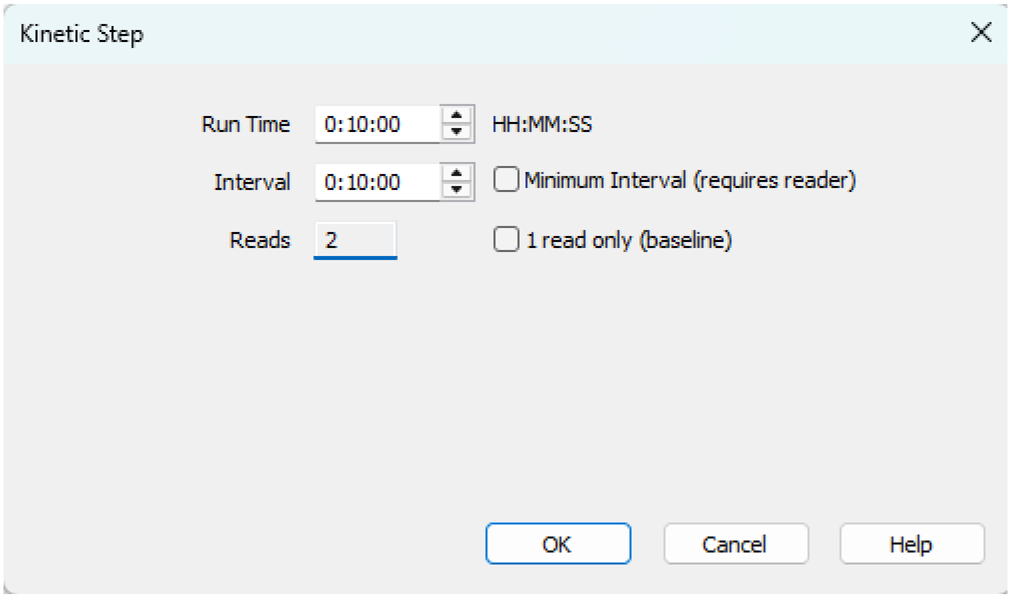


**Figure S5. LEMOS Protocol – Kinetic Step.**

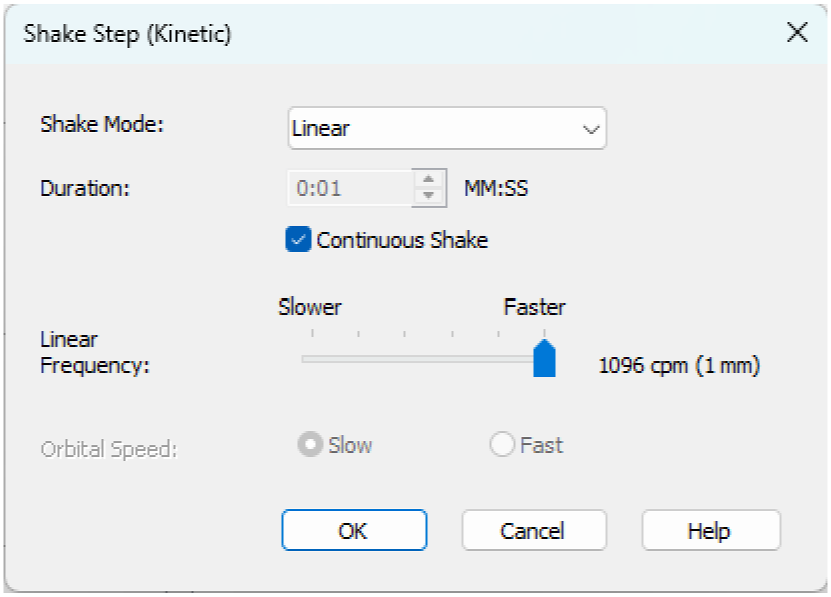


**Figure S6. LEMOS Protocol – Shaking Step.**

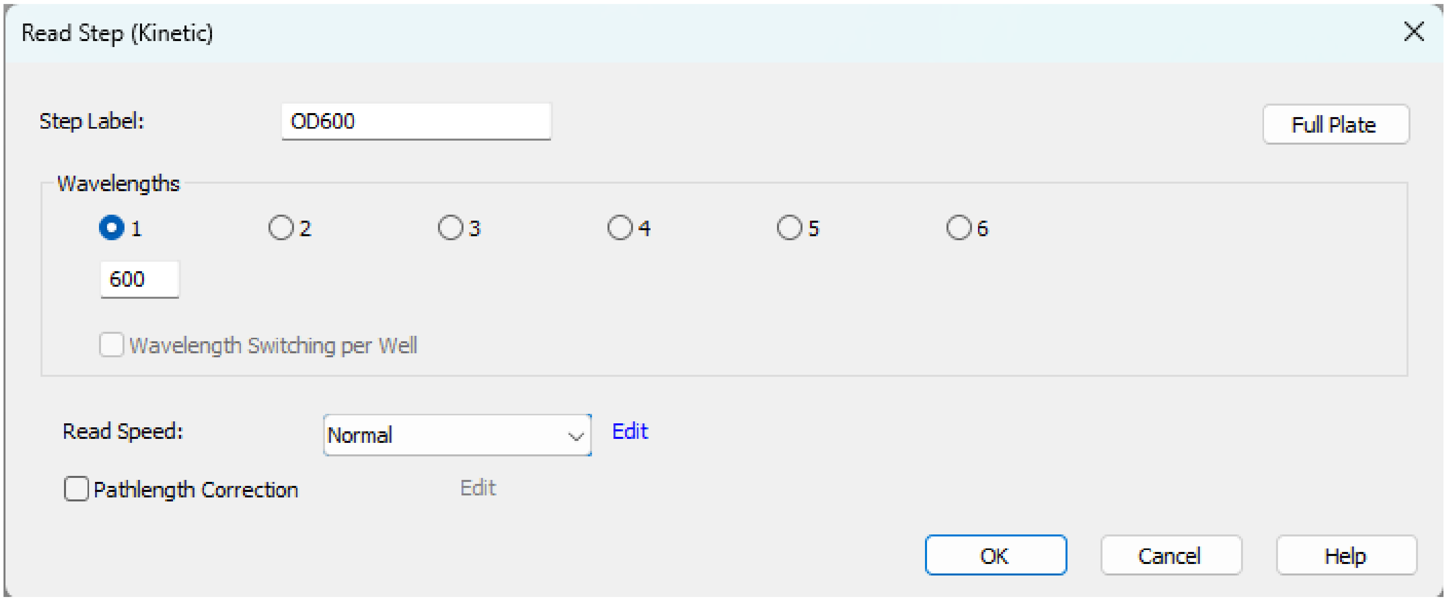


**Figure S7. LEMOS Protocol – Reading $OD_{600}$.**

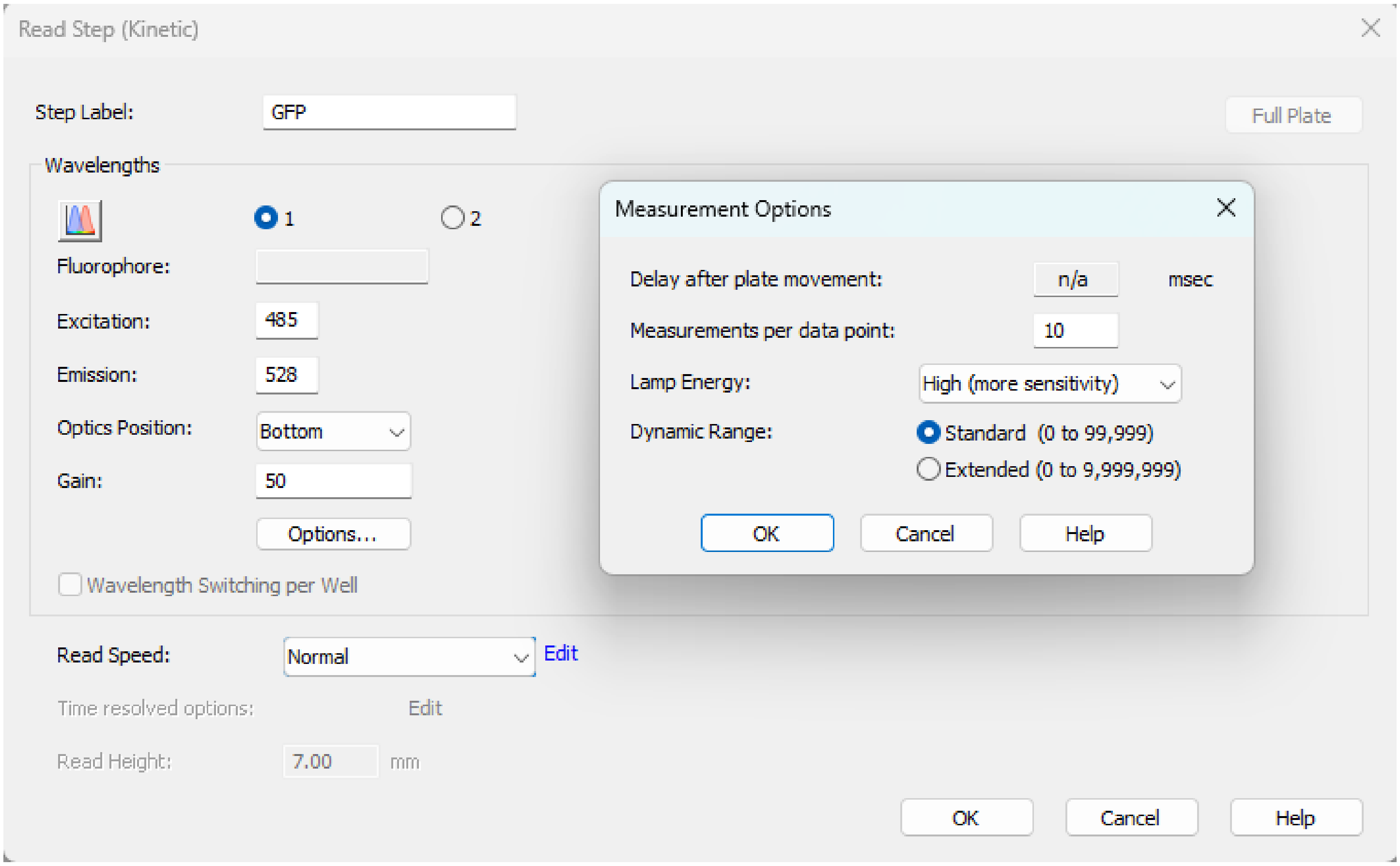


**Figure S8. LEMOS Protocol – Reading Fluorescence.**

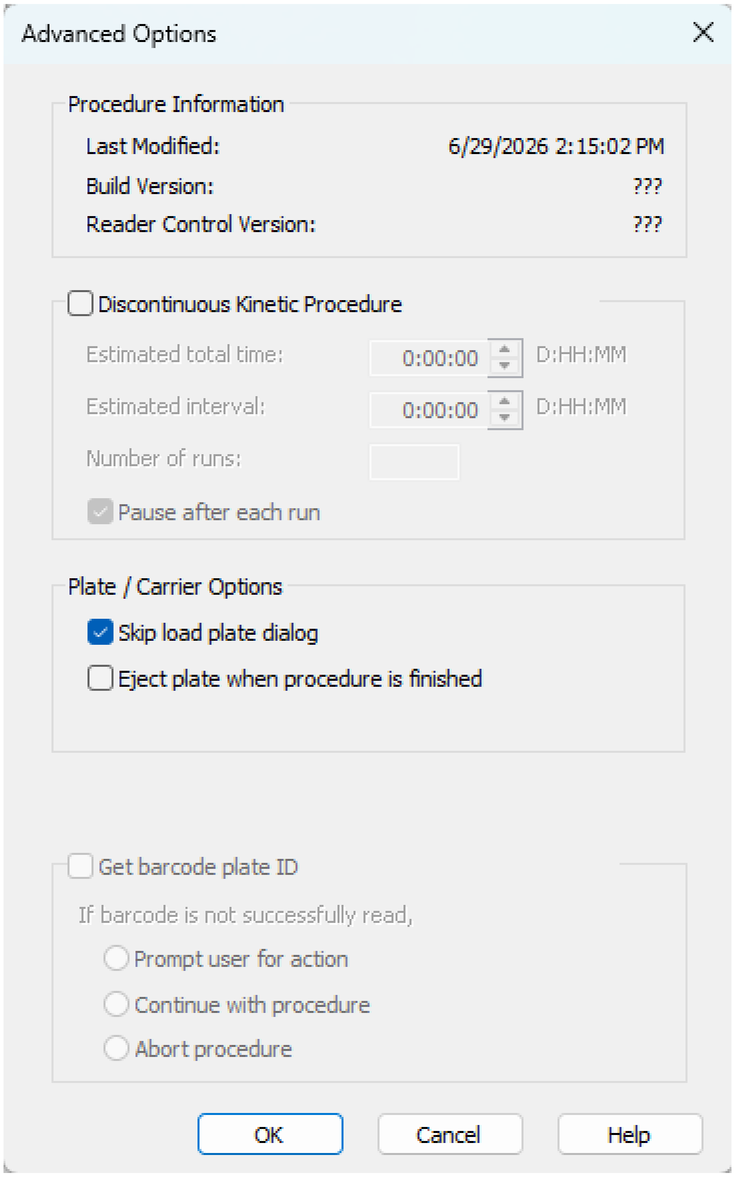


**Figure S9. LEMOS Protocol – Advanced Settings.**

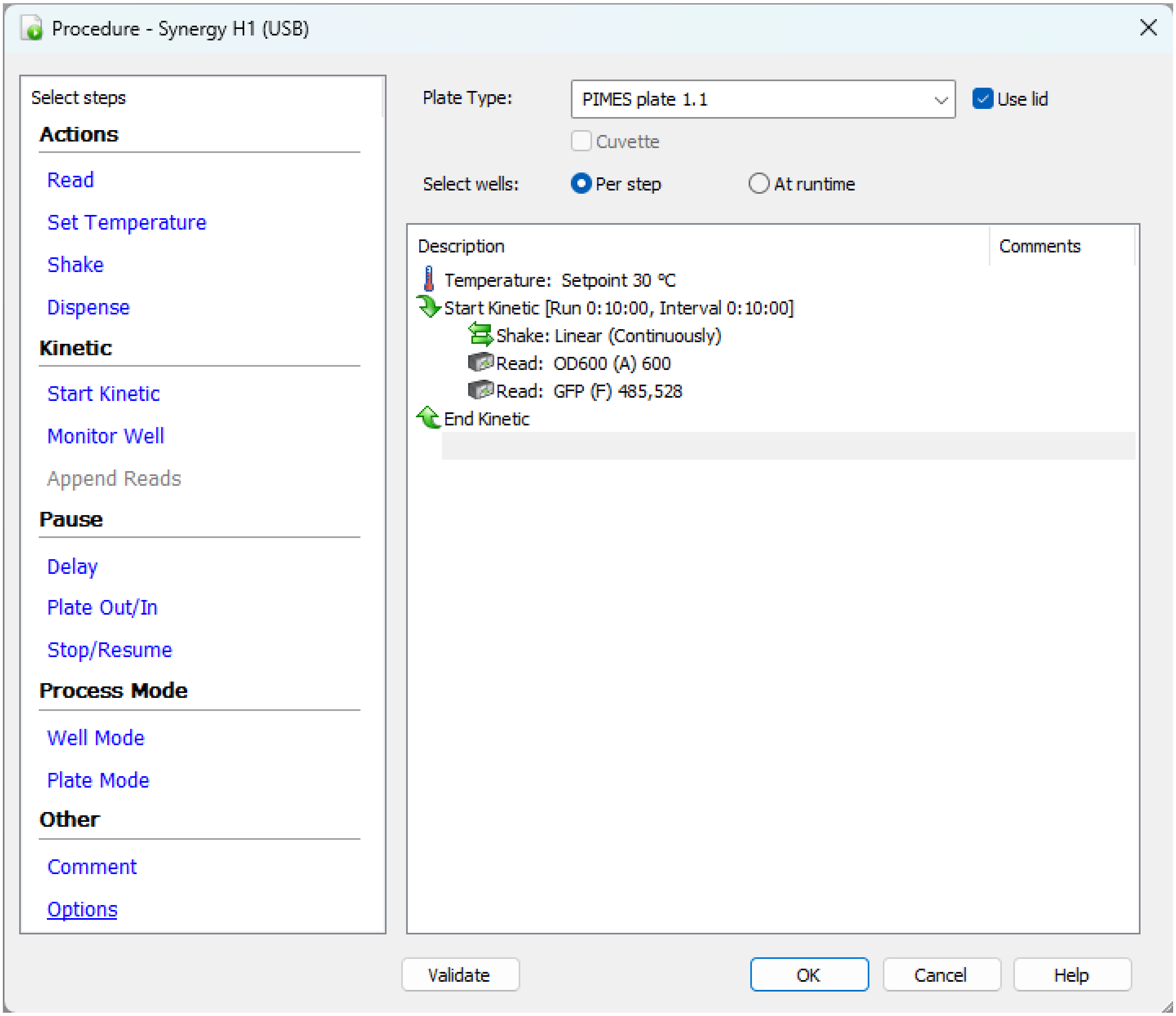


**Figure S10. LEMOS Protocol – Final Setup.**

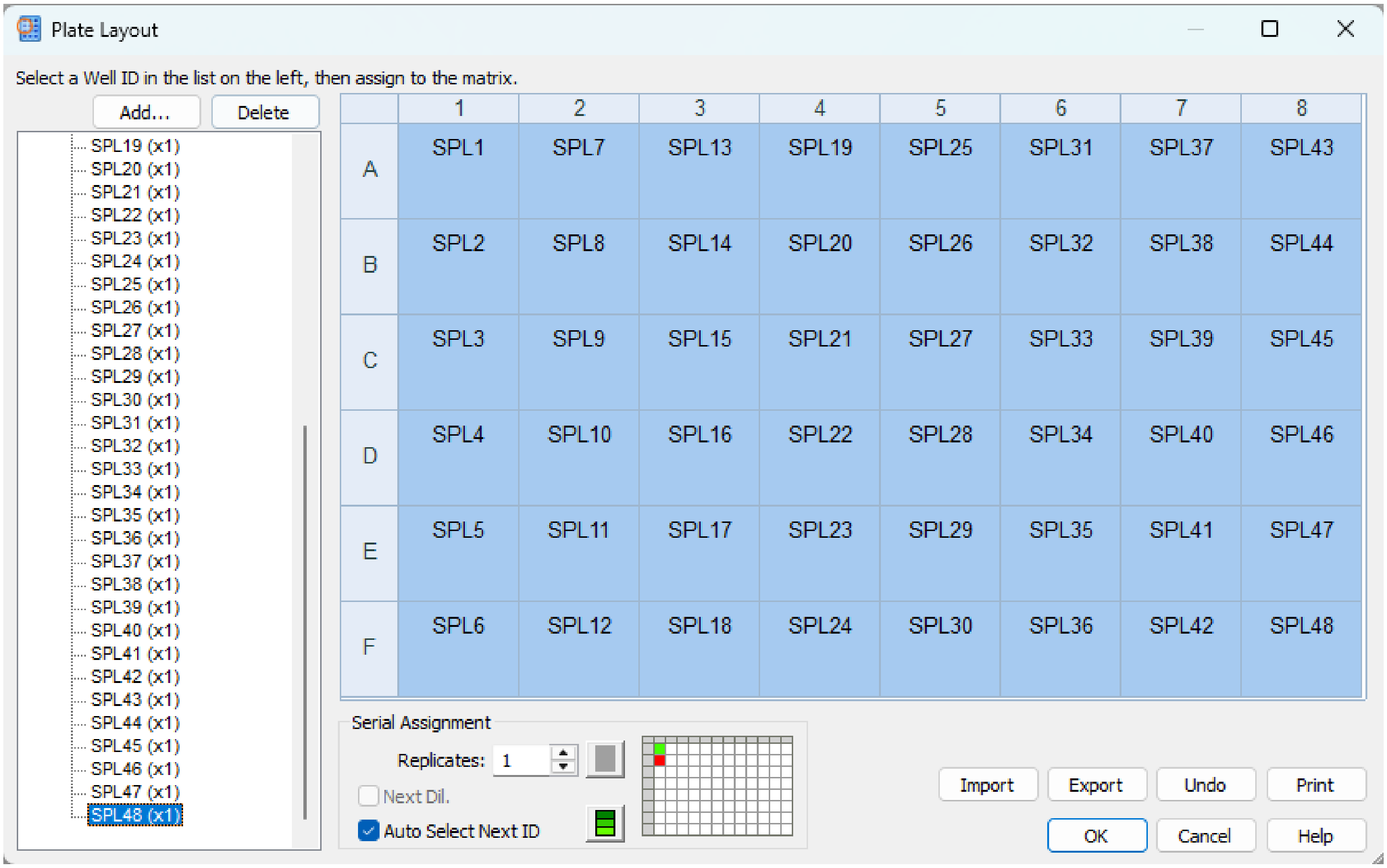


**Figure S11. LEMOS Plate Layout Settings.**

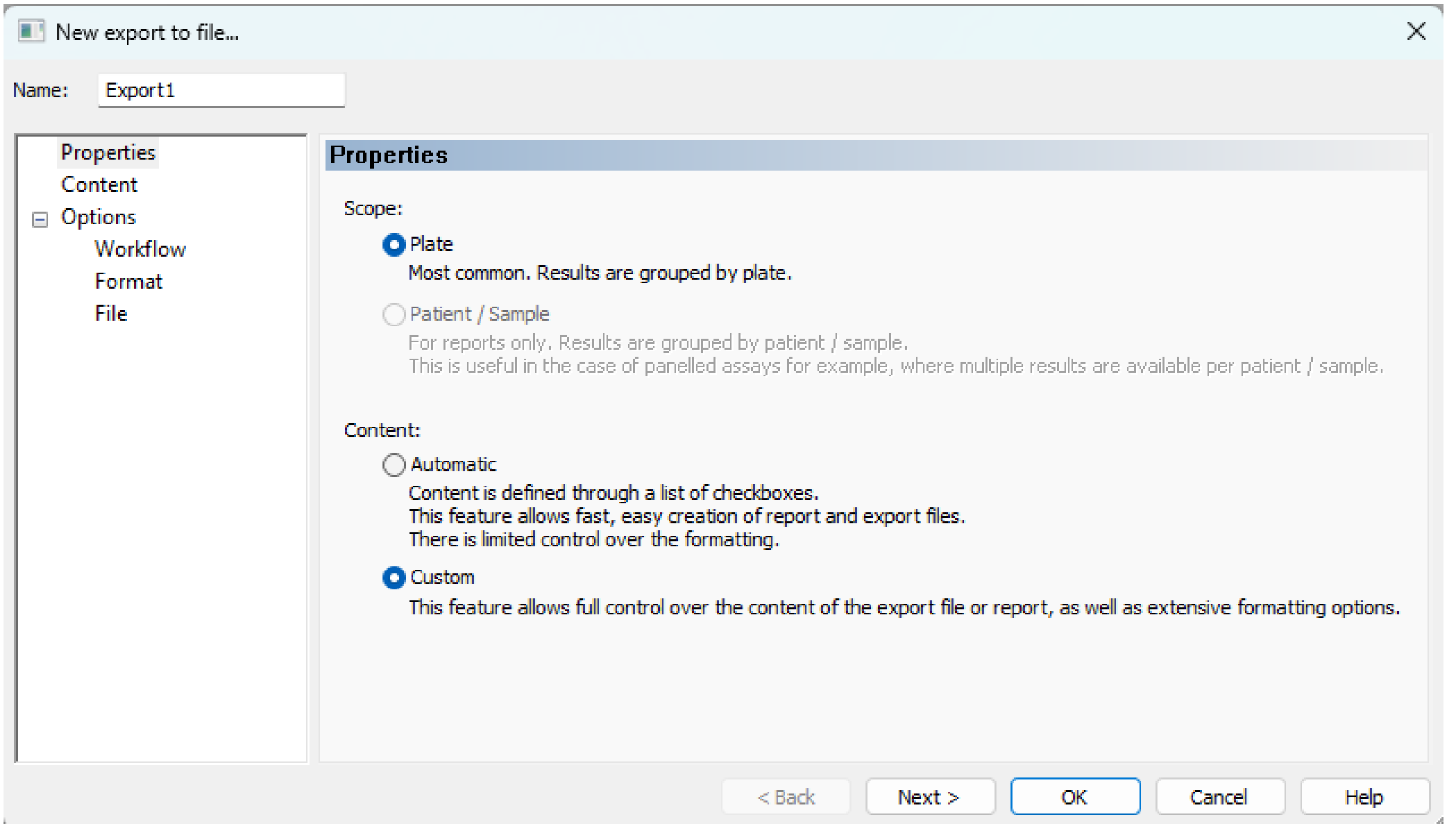


**Figure S12. LEMOS Export – Custom Content.**

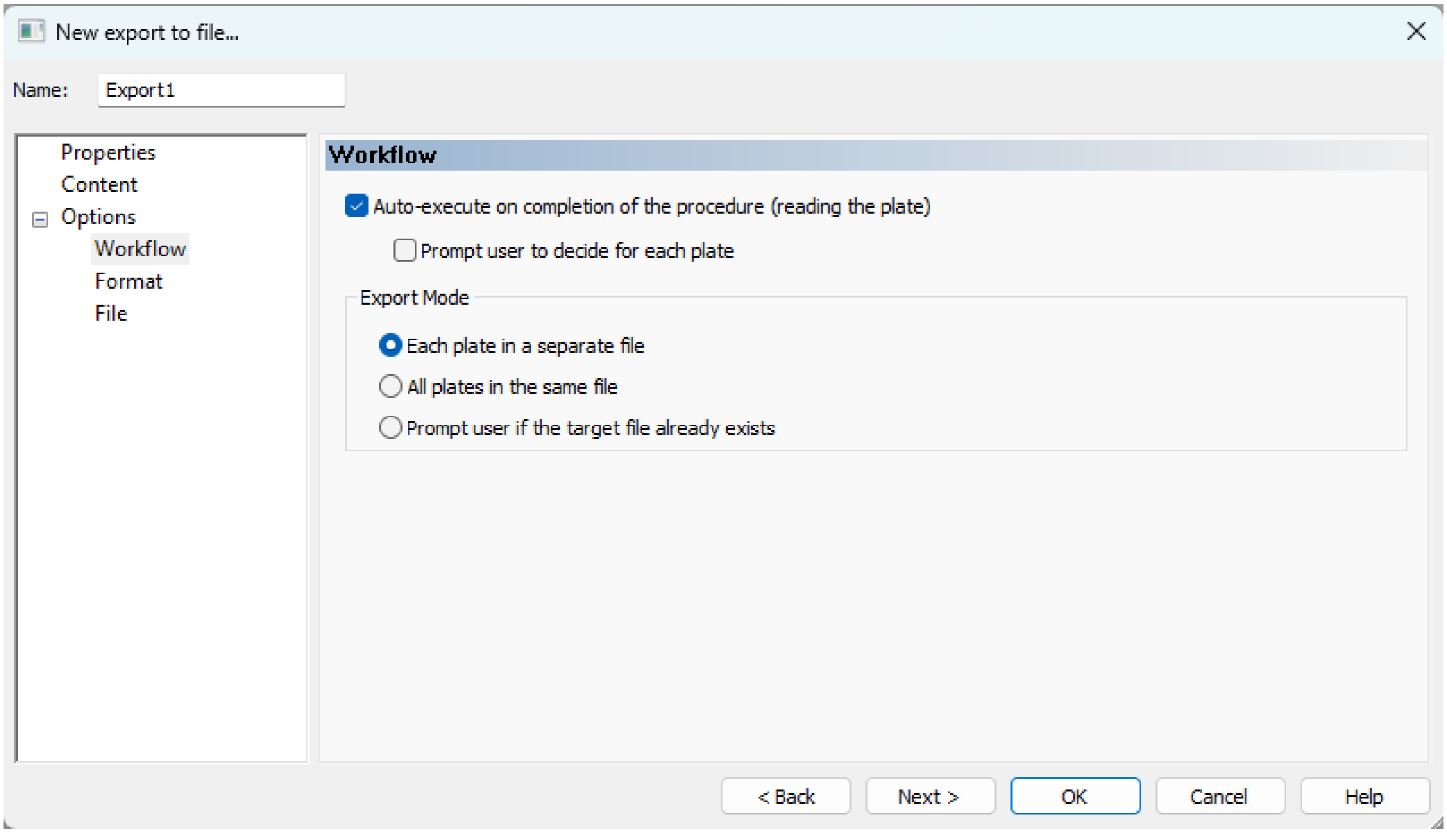


**Figure S13. LEMOS Export – Workflow.**

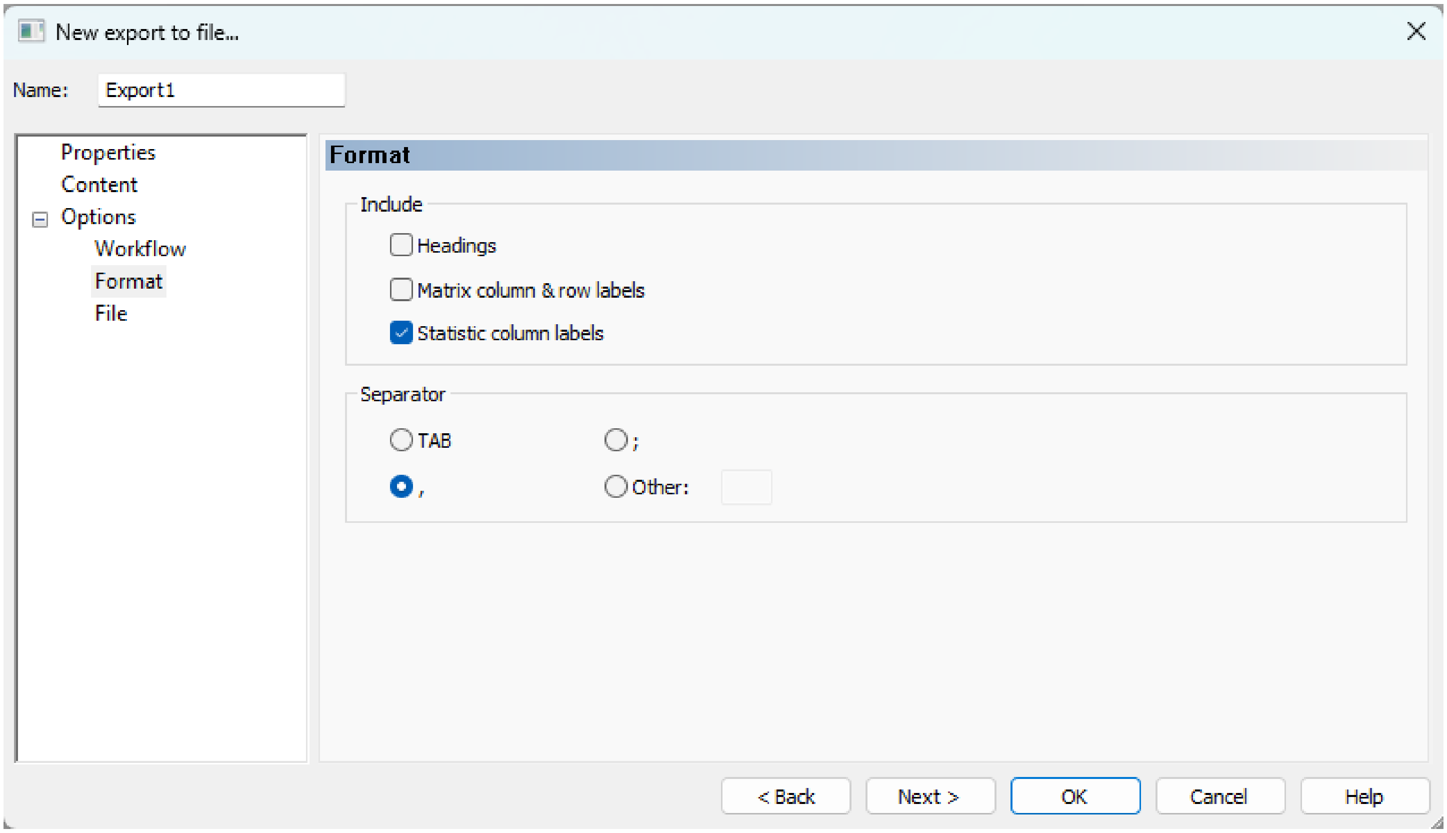


**Figure S14. LEMOS Export – Format.**

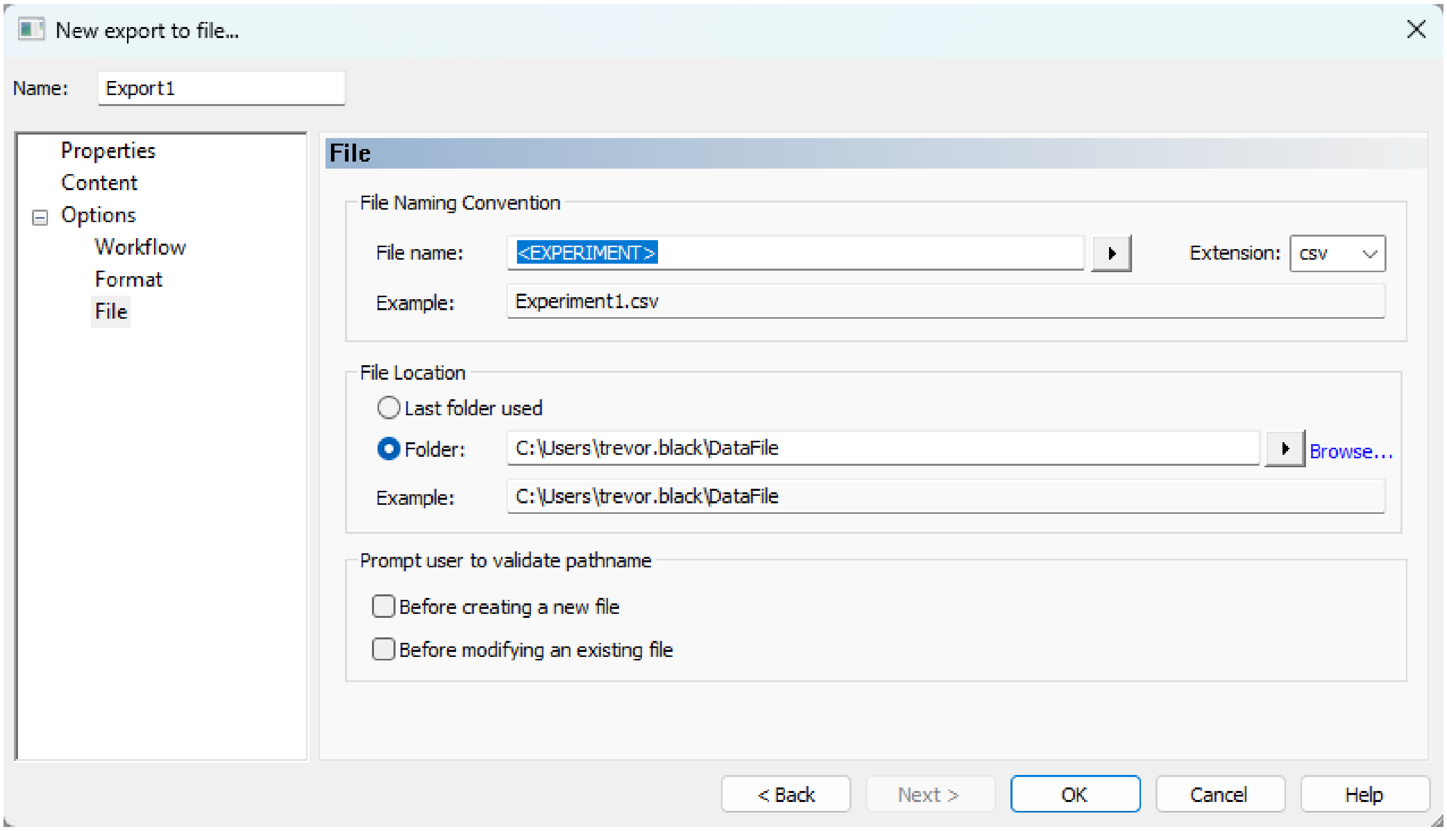


**Figure S15. LEMOS Export – File Naming.**

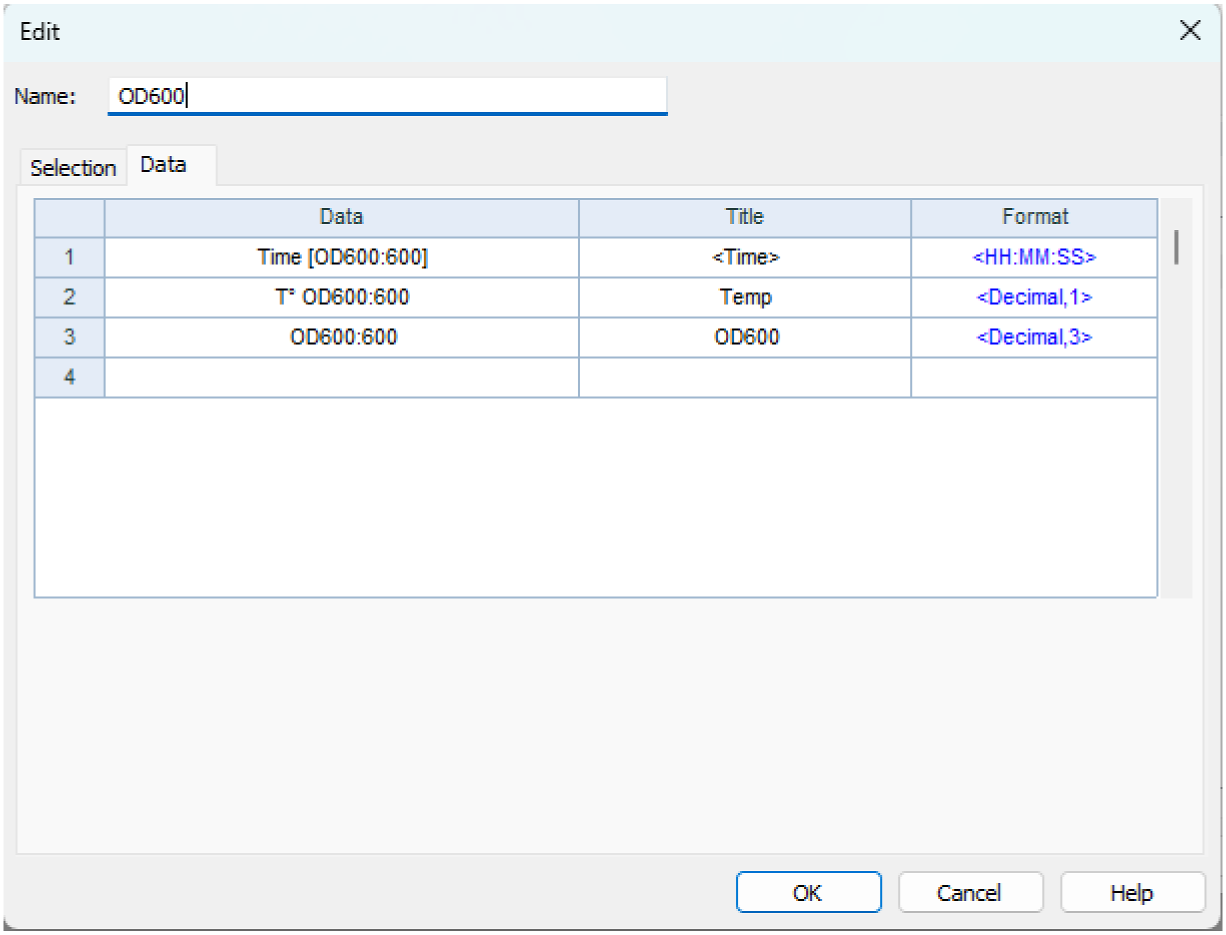


**Figure S16. LEMOS Export – Content Data $OD_{600}$.**

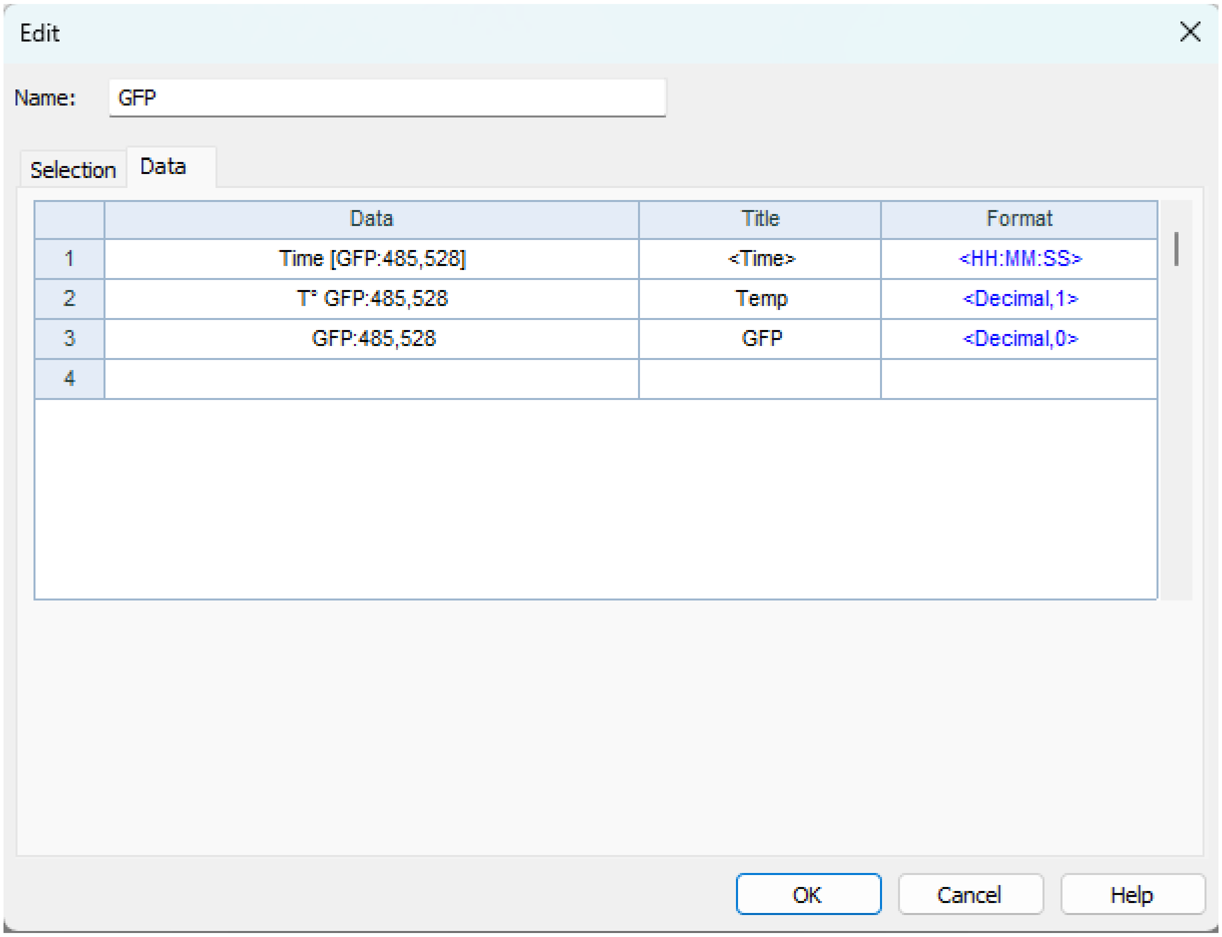


**Figure S17. LEMOS Export – Content Data GFP.**
